# Novel DNA methylation changes in mouse lungs associated with heavy smoking

**DOI:** 10.1101/2023.11.06.565206

**Authors:** Chinonye Doris Onuzulu, Samantha Lee, Sujata Basu, Jeannette Comte, Yan Hai, Nikho Hizon, Shivam Chadha, Maria Shenna Fauni, Andrew J. Halayko, Christopher D. Pascoe, Meaghan J. Jones

## Abstract

Smoking is a potent cause of asthma, chronic obstructive pulmonary disease (COPD) and many other health defects, and changes in DNA methylation (DNAm) have been identified as a potential link between smoking and these health outcomes. However, most links between smoking and DNAm have been made using blood and other easily accessible tissues in humans, while evidence from more directly affected tissues such as the lungs is greatly lacking. Here, we identified DNAm patterns which are altered by smoking directly in the lungs. We used a well-established mouse model to measure the effects of heavy smoking first on lung phenotype immediately after smoking and then after a period of smoking cessation. Next, we determined whether our mouse model could recapitulate previous DNAm patterns observed in smoking humans by measuring DNAm at a candidate gene responsive to cigarette smoke (CS), *Cyp1a1.* Finally we carried out epigenome-wide DNAm analyses using the newly released Illumina mouse methylation microarrays. Our results recapitulate some of the phenotypes and DNAm patterns observed in human studies but reveal 32 differentially methylated genes specific to the lungs which have not been previously associated with smoking. The affected genes are known to be involved in nicotine dependency, tumorigenesis and metastasis, immune cell dysfunction, lung function decline, and COPD. This research emphasizes the need to study CS-mediated DNAm signatures in directly affected tissues like the lungs, as that may be essential in understanding mechanisms underlying CS-mediated health outcomes.

## INTRODUCTION

Smoking is one of the leading causes of preventable mortality worldwide, and poses a great burden on the healthcare system, as it has been linked to the development of diseases like asthma, chronic obstructive pulmonary disease (COPD), bronchitis, obesity, and cancers^1–5^. Epigenetic mechanisms, particularly DNA methylation (DNAm), have been suggested as potential links between environmental exposures such as smoking and disease outcomes that might manifest later in life.

There is abundant evidence that smoking alters DNAm patterns in human peripheral blood^6–9^ and leucocyte subtypes^7^, lymphoblasts and pulmonary macrophages^10^, and epithelial oral cells^11^. In fact, there are reports that certain methylated CpGs are effective markers for predicting smoking status^12^ and lung cancer incidence^13^. However, most of these past studies associating smoking with DNAm alterations were done in accessible tissues, and while they are important in understanding how smoking affects the body as a whole, evidence from more proximal tissues such as lungs is severely lacking^14^. Even though some DNAm patterns overlap between tissues, it is important to measure DNAm in the specific tissue of interest, as DNAm is cell type and tissue-specific^15–17^. When directly comparing tissues from the same patients, many CpGs are differentially methylated between lung tissue and blood^18^. One recent study measured epigenome-wide DNAm in current smokers and reported that only a third of the significantly altered CpGs which they originally identified in whole blood were also altered in the lungs of current compared to never-smokers ^14^. Another study compared smoking-induced DNAm changes in blood versus buccal cells and reported that DNAm sites from buccal cells outperformed those identified in blood cells in discrimination of 14 of 15 epithelial cancer types^19^. The latter study showed that tissues directly exposed to cigarette smoke (CS) such as lungs, buccal cells and saliva may be more accurate in detecting direct effects of smoking on DNAm. It can be difficult to access these tissues in humans, so model organisms are often employed.

Past research using animal models to measure the effects of smoking on DNAm in the lungs is limited as well^14,20,21^. One study which measured the effect of four weeks of smoking on lung DNAm using capture based bisulfite sequencing found that smoking altered DNAm patterns at numerous genes, most of which were involved in inflammation and in inflammatory injury in COPD^21^. As this is one of very few, the need for more research to add to the existing ones cannot be overemphasized.

The DNAm microarray is a high-throughput method for measuring epigenome-wide DNAm at a lower cost and less labour intensive than bisulfite sequencing. Sequentially increasing scales of the Infinium microarrays have been developed for human samples, but no such solution was available for use on mouse samples until recently, with the release of the Illumina Infinium mouse microarrays^22^. In this study, we have used an animal model of heavy smoking and measured the effects of smoking on lung function, gene expression and DNAm of a candidate gene immediately after smoking and 15 weeks after smoking cessation, and using the new mouse methylation array, we measured epigenome-wide DNAm immediately after smoking. Our model successfully recapitulated some of the lung phenotypes and DNAm signatures observed in humans. It also reveals many novel sites in the lungs which are differentially methylated due to smoking, many of them involved in nicotine dependence, altered immune response and development of malignancies, lung function decline and COPD, cardiovascular defects and neurodegenerative disorders. This study is an essential step in understanding mechanisms underlying CS-induced health alterations, and therefore would help in developing therapies to mitigate the effects of smoking.

## METHODS

### Animals and smoking model

This experiment was approved by the Animal Research Ethics and Compliance Committee of the University of Manitoba. Adult Balb/C mice (Charles River Laboratories, Massachusetts, United States) were given standard laboratory chow and clean water *ad libitum* and housed, four mice of a single sex per cage (except where noted), in individually ventilated cages in a 12-hour light/dark cycle throughout the duration of this experiment.

The full details of the smoke exposure paradigm have been published previously^23^. Briefly, 8-week-old adult female Balb/C mice were separated into 2 groups: 16 control mice and 16 smoke-exposed mice. Treatments were for a total period of 9 weeks, beginning 3 weeks prior to mating (at which point mice were singly housed), continuing throughout pregnancy and lasting until 3 weeks after birth of offspring. Control mice were moved to a clean cage and exposed to room air, while CS mice were exposed to whole body 1R6F research cigarettes (University of Kentucky, Lexington, KY), twice daily using the SCIREQ InExpose smoking robot (SCIREQ, Montreal, QC, Canada). The results reported in this study are a part of a larger experiment to investigate the effects of maternal smoking on offspring^23^. Therefore, the ‘dams’ referred to from this point forward were all pregnant in line with that experiment. Dams were weighed weekly and 48 to 72 hours after the last smoke exposure (referred to hereafter as “immediately after smoking”), a subset of dams underwent lung function testing followed by tissue collection. The remaining dams underwent lung function testing and tissue collection 15 to 16 weeks after smoking cessation (referred to hereafter as “15 weeks after smoking cessation”), resulting in data from 2 timepoints: immediately after 9 weeks of smoking and 15 weeks after smoking cessation (Sample and group information can be found in Supplementary Table S1).

### Measurement of maternal cotinine

To verify delivery of CS, cotinine levels were measured in plasma immediately after smoking, and again 15 weeks later in a subset of dams. Cotinine was measured using a commercially available ELISA kit (CalBiotech, Spring Valley, CA), according to the manufacturer’s instructions, and the concentration of cotinine in each sample was measured at 450 nm.

### Lung function measurement and Bronchoalveolar lavage fluid collection

Mice were anesthetized with sodium pentobarbital and lung function measured using the SCREQ Flexivent small animal ventilator (SCIREQ Inc., Montreal, QC, Canada) as described previously^24,25^. Four lung function metrics: Total airway resistance (Rrs), Newtonian resistance (Rn), tissue resistance (G) and elastance (H) were assessed, first at baseline after injection of nebulized saline into the lungs, and then after introduction of increasing concentrations of nebulized methacholine (3 to 50 mg/mL).

Once lung function measurements were complete, mouse lungs were washed twice with 1 mL of phosphate buffer saline (PBS) per wash, introduced via tracheal cannulation. The bronchoalveolar lavage fluid (BALF) obtained was centrifuged at 4°C at 1200 rpm for 10 min, and the supernatants were stored at −20°C to be used for future analysis. Cell pellets were resuspended in 1 ml of PBS for total cell counts using a hemocytometer. Differential cell counts of macrophages, eosinophils and lymphocytes were performed by first pipetting 100ul of PBS-resuspended cell pellets onto glass slides using cytospin columns, staining with a modified Wright-Giemsa stain (Hema 3 Stat Pack), and then counting cells using a Carl Zeiss Axio Observer ZI microscope.

### Tissue collection, DNA/RNA isolation

We collected whole blood from dams immediately after 9 weeks of CS exposure and 15 weeks after smoking cessation. Blood obtained from the severed abdominal aorta was immediately pipetted into EDTA-coated tubes and centrifuged at 4000 rpm for 15 min to obtain plasma. Following separation, blood cell pellets and plasma were snap-frozen in dry ice and stored at −80°C. Next, left, middle, superior, inferior and postcaval lung lobes were collected into separate tubes and snap-frozen in dry ice.

To prepare for simultaneous DNA and RNA extraction using the Invitrogen DNA and RNA isolation kits, we homogenized whole left lungs in Qiagen RLT Plus buffer using the Qiagen Tissue Lyser II. We also extracted DNA from blood cell pellets using the Qiagen DNAeasy Blood and Tissue kit and quantified all extracted DNA and RNA using a NanoDrop spectrophotometer (NanoDrop Technologies, USA).

### Selection of candidate genes to measure DNAm in mice

As our mouse model was novel, we needed to confirm that we could use it to recapitulate some of the DNAm signatures previously reported in humans exposed to CS. We decided to measure DNAm at candidate genes first on a small scale and then proceed to epigenome-wide DNAm measurements. Complete details of how we selected our 2 candidate genes have been published previously^23^. Briefly, we selected 2 CpGs which were the most responsive to *in utero* CS in newborn umbilical cord blood samples as reported in a meta-analysis conducted across human cohorts, and had also been associated with exposure to CS in adulthood^26^. The two chosen human CpGs were *AHRR* (cg05575921) and *CYP1A1* (cg22549041). We then aligned these sequences against the GRCm38/mm10 *Mus musculus* genome assembly using muscle^27^ alignment on R studio version 3.6. This produced mouse CpGs which exactly aligned with or were closest in position to the human CpGs selected from the meta-analysis. The two mouse CpGs were at chr13:74260517 for *Ahrr* and chr9:57696231 for *Cyp1a1* (GRCm38/mm10). It is essential to note that this region of *Ahrr* is not conserved between humans and mice, and while we selected the closest mouse CpG, it may not be comparable to the human position.

### Gene expression measurement

200 ng of lung mRNA was converted to cDNA using Maxima cDNA synthesis Kit (Thermo Fisher Scientific, Inc.) following manufacturer’s protocol. We measured relative expression of *Ahrr* and *Cyp1a1* in dam lungs using quantitative real-time RT-PCR (qPCR) performed on the QuantStudio 3 Real-Time PCR System (Thermo Fisher Scientific, Waltham, MA). *Ahrr* and *Cyp1a1* expression was quantified using the ΔΔ^Cq^ method^28,29^ after normalizing against mean β*-actin* and *Eif2a* levels in the same sample. Samples were run in duplicates under cycling conditions that were recommended by the manufacturer, and the primer sequences used have been published previously^23^.

### Measurement of DNAm at the candidate genes, *Ahrr* and *Cyp1a1*

We measured DNAm at the two chosen mouse candidate genes, *Ahrr* and *Cyp1a1,* via pyrosequencing. Briefly, we performed bisulfite conversion (Zymo Research) on 500 ng of DNA isolated from the left lungs or blood to generate bisulfite-converted DNA (bcDNA), following the manufacturer’s protocol. Converted DNA was amplified using primers and conditions published previously^23^. DNAm at candidate genes was measured in duplicates using the Qiagen Q48 pyrosequencer, alongside mouse control DNA of increasing methylation concentrations: 0%, 25%, 50%, 75%, and 100%.

### Measurement of epigenome-wide DNAm in dam lung autosomes

We used the Illumina Infinium Mouse Methylation BeadChip (Illumina, San Diego, CA, USA) to measure DNAm across the whole mouse epigenome in accordance to the manufacturer’s protocol. Briefly, 750 ng of genomic DNA was bisulfite converted as described above. Next, bisulfite converted samples were randomized, amplified, fragmented, hybridized onto the array chip and scanned according to the standard protocol.

We exported IDAT files were exported into R and performed preprocessing using the SeSAMe package^30^. Preprocessing steps included removal of 18,920 of 296,070 probes mapping to the X, Y and mitochondrial chromosomes, calculation of detection *p* values per probe using pOOBAH^30^, background subtraction using noob^31^, dye bias correction using the dyeBiasCorrTypeINorm function in the SeSAMe package and filtering out probes with detection *p* > 0.05. Next, signal intensities of the remaining 208,606 probes were quantified as β-values and used in downstream analyses.

### Measurement of epigenome-wide DNAm in dam lung X and Y chromosomes

We measured the effect of smoking for 9 weeks on DNAm in Illumina mouse microarray probes mapping to the female X chromosomes separately. Similar to autosomal probes, preprocessing was conducted using the SeSAMe package in R^30^. We performed the preprocessing steps highlighted above on 15,117 of 296,070 probes mapping to the female X chromosome, and were left with 11,529 probes used in downstream analyses.

### Statistical analyses

Statistical analysis were performed in RStudio (versions 3.6.1 and 4.2). To analyze candidate gene DNAm data, we averaged DNAm at each CpG across duplicates and measured between-group differences using student t-tests. We used one-way ANOVA to analyze lung function and differential cell count data, followed by student t-tests for multiple comparisons between groups at each methacholine dose. We considered *P* values < 0.05 as significant.

To analyze epigenome-wide DNAm microarray data, we first performed chip, run, row and column batch correction on β values using ComBat^32^, included in the SVA package^33^. Next, we used the SVA package to capture technical covariates to be included in our linear regression model, and included the recommended 1 surrogate variable in our model. Next, we used multivariable linear regression contained within the limma R package to measure differential DNAm on the β values. Since our study design has a very small sample size (N = 3 control, N = 3 CS) and our *p* values would therefore not accurately reflect very small differences^34^ in DNAm, we considered CpGs as significant at *p* < 10e-3 and an effect size > 0.05. We produced figures in R using the *ggplot2* and *Gviz* packages.

### Gene Ontology

To obtain biological significance from identified differentially methylated probes, we used the ClusterProfiler R package^35^ (v3.16.1) for gene enrichment analysis. We considered pathways with FDR < 0.05 to be significantly enriched.

## RESULTS

The research presented here is one part of a larger animal experiment to investigate the effects of prenatal CS exposure on offspring. Therefore, the ‘dams’ we refer to is in reference the fact that these female mice were pregnant at some point. We exposed Balb/C dams (N = 16 controls and N = 16 CS-exposed) to whole-body CS for a total period of 9 weeks, starting 3 weeks before mating, and lasting throughout pregnancy until 3 weeks after the birth of their pups (Figure 1A). CS-exposed dams tolerated smoking, with no losses or adverse events during the course of the experiments. We performed lung function and collected tissues for further analyses at 2 timepoints: 48-72 hours after the last day of 9 weeks of smoking (referred to hereafter as “immediately after smoking”), and 15-16 weeks after smoking cessation (referred to hereafter as “15 weeks after smoking cessation”) (Figure 1A, Supplementary Table S1).

**Figure 1:**
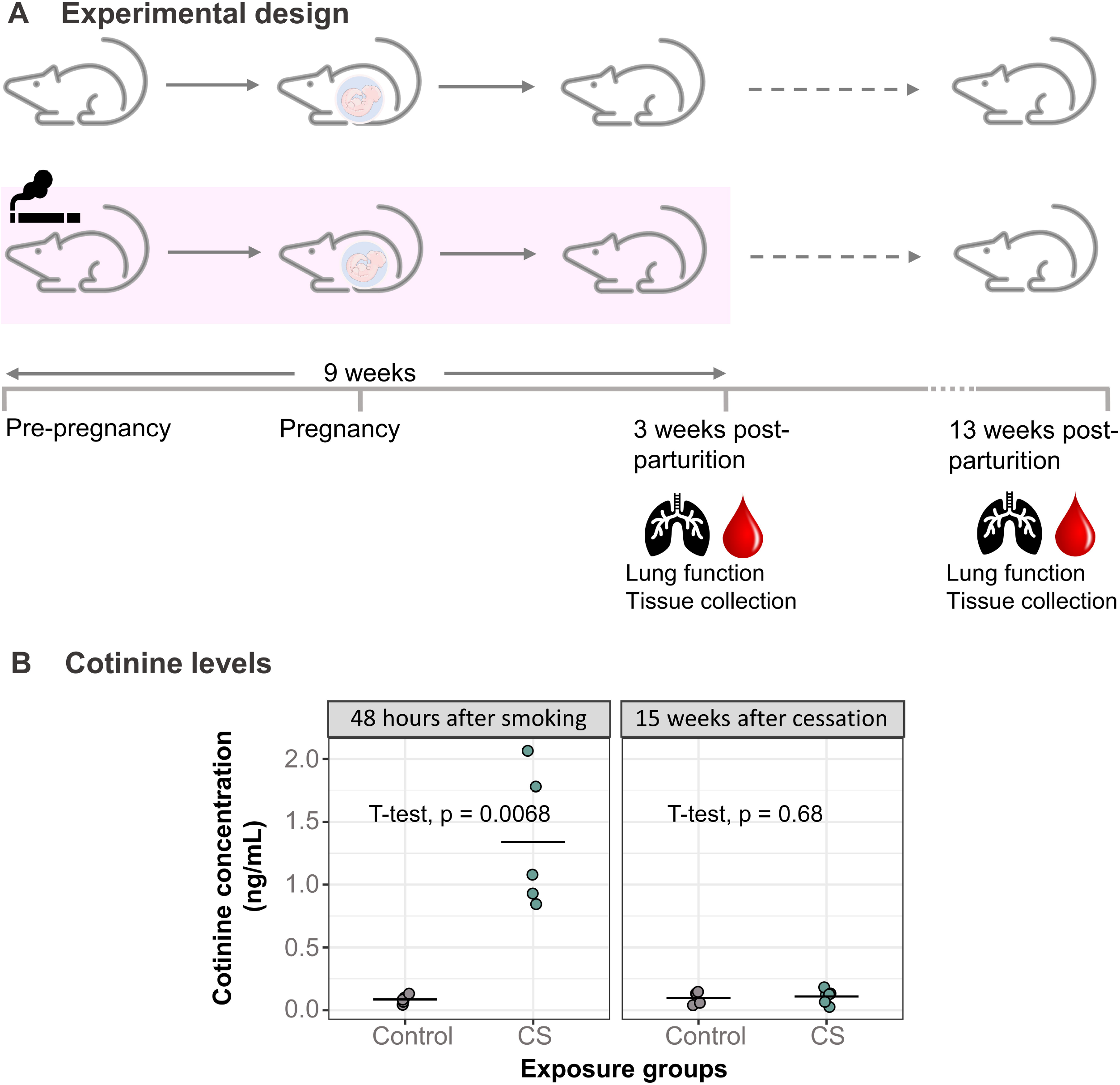
Development of a mouse model to study the effects of heavy smoking. (A) Adult female mice (N = 16 control, N = 16 CS) were exposed to heavy doses of whole-body CS for 9 weeks, from a pre-pregnancy period of 3 weeks and ending 3 weeks after birth of offspring. Following lung function measurements, tissues were collected from dams immediately after the last day of 9 weeks of smoke exposure and then 15 weeks after smoking cessation. (B) Dam cotinine levels measured in plasma 48 hours and 15 weeks after smoking cessation. N = 5 per group. Differences in plasma cotinine were analyzed using student t-tests.

To verify absorption of CS components, we measured cotinine levels in plasma collected at these 2 different timepoints (Figure 1B). Dams which were exposed to CS had 15 times more elevated plasma cotinine levels (Mean = 1.34, SD = 0.55) immediately after smoking cessation compared to control dams (Mean = 0.09, SD = 0.03) (Figure 1B, *p* = 0.0068). The relatively low levels of cotinine observed overall in the smoke-exposed dams can be attributed to the collection of plasma from dams approximately 48 hours after smoking, with the reported half-life of cotinine being about 24 hours^36,37^. Cotinine levels measured 15 weeks after CS exposure showed no significant difference between control and smoking dams (Figure 1B).

### Lung phenotype immediately after 9 weeks of smoking and 15 weeks after smoking cessation

Several studies have shown that personal smoking alters lung function and induces airway hyperresponsiveness^38–40^. Therefore, we assessed the effects of heavy smoking on the lung phenotype by measuring lung function and immune cell infiltration.

We found that direct CS exposure did not alter dam lung function either at baseline or after methacholine challenge after 9 weeks of smoking, though tissue elastance was lower but not statistically significant in CS exposed than unexposed dams (Figure 2E-2H and Figure 3A-3D). Thirteen weeks after smoking cessation, baseline lung function was also not altered (Figure 2E-2H), but we observed significant differences between CS exposed and unexposed animals in tissue elastance, tissue resistance and total lung resistance values (Figure 3E-3H). Smoking for 9 weeks also caused a 4-fold and 8-fold increase in eosinophils and lymphocytes respectively immediately after smoking, compared to controls (Figure 2A-2D). The immune cell infiltration differences, however, did not persist until 15 weeks after smoking cessation (Figure 2A-2D). Also of note is that total cell count and macrophage cell count were higher in both exposure groups at the immediate post-smoking timepoint, when the dams were 3 weeks post-parturition and still lactating, than 15 weeks later.

**Figure 2:**
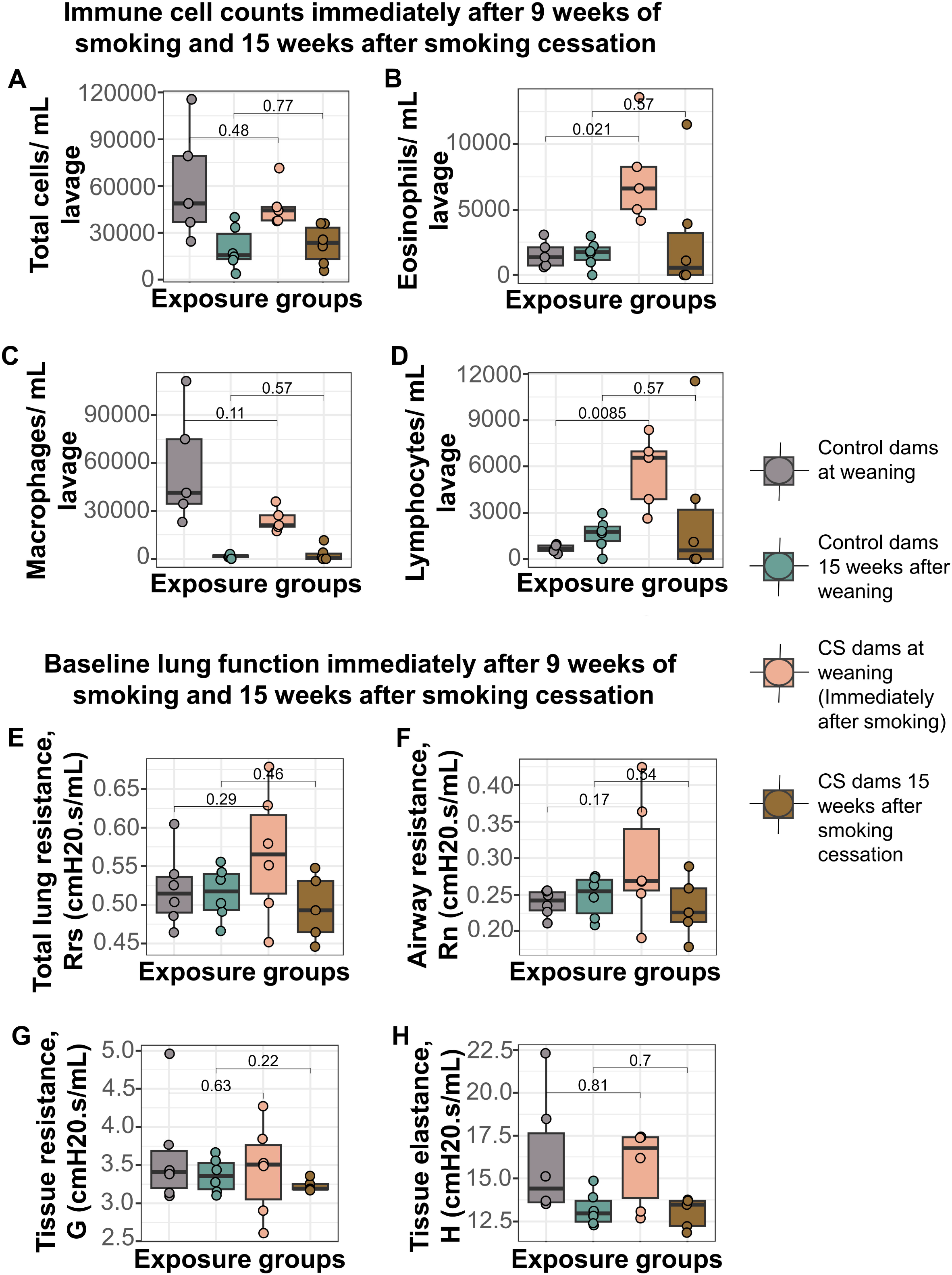
Smoke exposure causes immediate but transient changes to immune cell infiltration into dam lungs, but does not alter dam baseline lung function over time. (A) Total immune cells per mL lavage immediately after smoking and 15 weeks after smoking cessation. (B) Eosinophils per mL lavage in the CS-exposed dams (Mean = 7528, SD = 3736) was over 4 times more elevated than control dams (Mean = 1569, SD = 1029) immediately after smoking, but normalized to controls 15 weeks after smoking cessation. (C) Macrophages per mL lavage immediately after smoking. (D) Lymphocytes per mL lavage in the CS-exposed dams (Mean = 5680, SD = 2357) was over 8 times elevated compared to control dams (Mean = 649, SD = 256) immediately after smoking, but normalized to controls 15 weeks after smoking cessation. (E) Total lung resistance at baseline. (F) Airway resistance at baseline. (G) Tissue resistance at baseline. (H) Alveolar elastance at baseline. N = 4-6 per group. Differential cell counts were normalized to lavage volume. Lung function values were measured using 90^th^ percentile values after injection of saline into the lungs. Two-group comparisons were conducted using a student t-test and *p <* 0.05 was considered significant.

**Figure 3:**
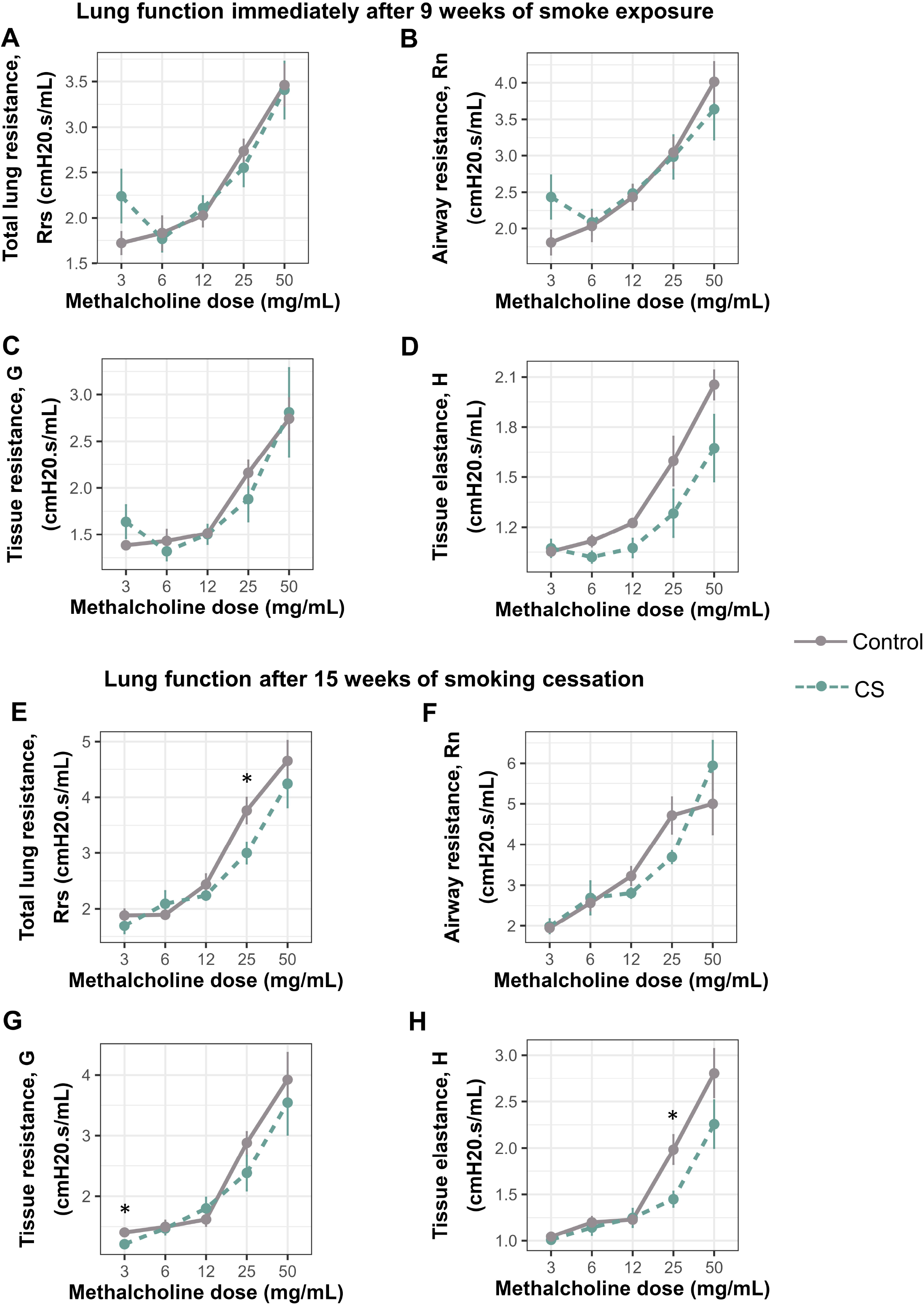
Smoke exposure alters methacholine responsiveness in mouse lungs up to 15 weeks after smoking cessation. (A) Total lung resistance immediately after smoking. (B) Airway resistance immediately after smoking. (C) Tissue resistance immediately after smoking. (D) Alveolar elastance immediately after smoking. (E) Total lung resistance after 15 weeks of smoking cessation. (F) Airway resistance after 15 weeks of smoking cessation. (G) Tissue resistance after 15 weeks of smoking cessation. (H) Alveolar elastance after 15 weeks of smoking cessation. N = 5-6 per group. Lung function values were measured using 90^th^ percentile values upon administration of increasing doses of methacholine. Data was analyzed using one-way ANOVA, followed by multiple comparisons at each methacholine dose where significant. \**p <* 0.05 in control vs smoke-exposed groups.

Together, this data showed that heavy smoking for 9 weeks alters lung phenotype by increasing immune cell infiltration into the lungs immediately after smoking, accompanied by persistent decline in lung function even after smoking cessation.

### Candidate gene DNAm and expression immediately after 9 weeks of smoking and 15 weeks after smoking cessation

To determine whether CS-induced DNAm changes observed in humans are recapitulated in our mouse model, we identified candidate genes to assess initially via pyrosequencing. We first measured DNAm and expression at *Cyp1a1* immediately after smoking and 15 weeks after cessation. We found an increase in *Cyp1a1* DNAm in the blood of CS-exposed dams 15 weeks after smoking cessation compared to controls, though this increase was not statistically significant (Figure 4A). Chronic exposure to CS for 9 weeks led to a significant decrease in *Cyp1a1* DNAm in the left lungs of CS-exposed dams immediately after smoking (Figure 4C, Mean = 11.20, SD = 1.05), compared to controls (Figure 4A, Mean = 19.20, SD = 0.80). Interestingly, *Cyp1a1* DNAm remained significantly decreased in the lungs of smoking dams (Figure 4D, Mean = 15.00, SD = 1.07) compared to controls (Figure 4B, Mean = 17.60, SD = 0.58) till 15 weeks after smoking cessation, though the magnitude had reduced slightly (Figure 4). *Cyp1a1* expression in the lungs of CS-exposed dams was slightly but not significantly elevated 15 weeks after smoking cessation (Figure 4B).

**Figure 4:**
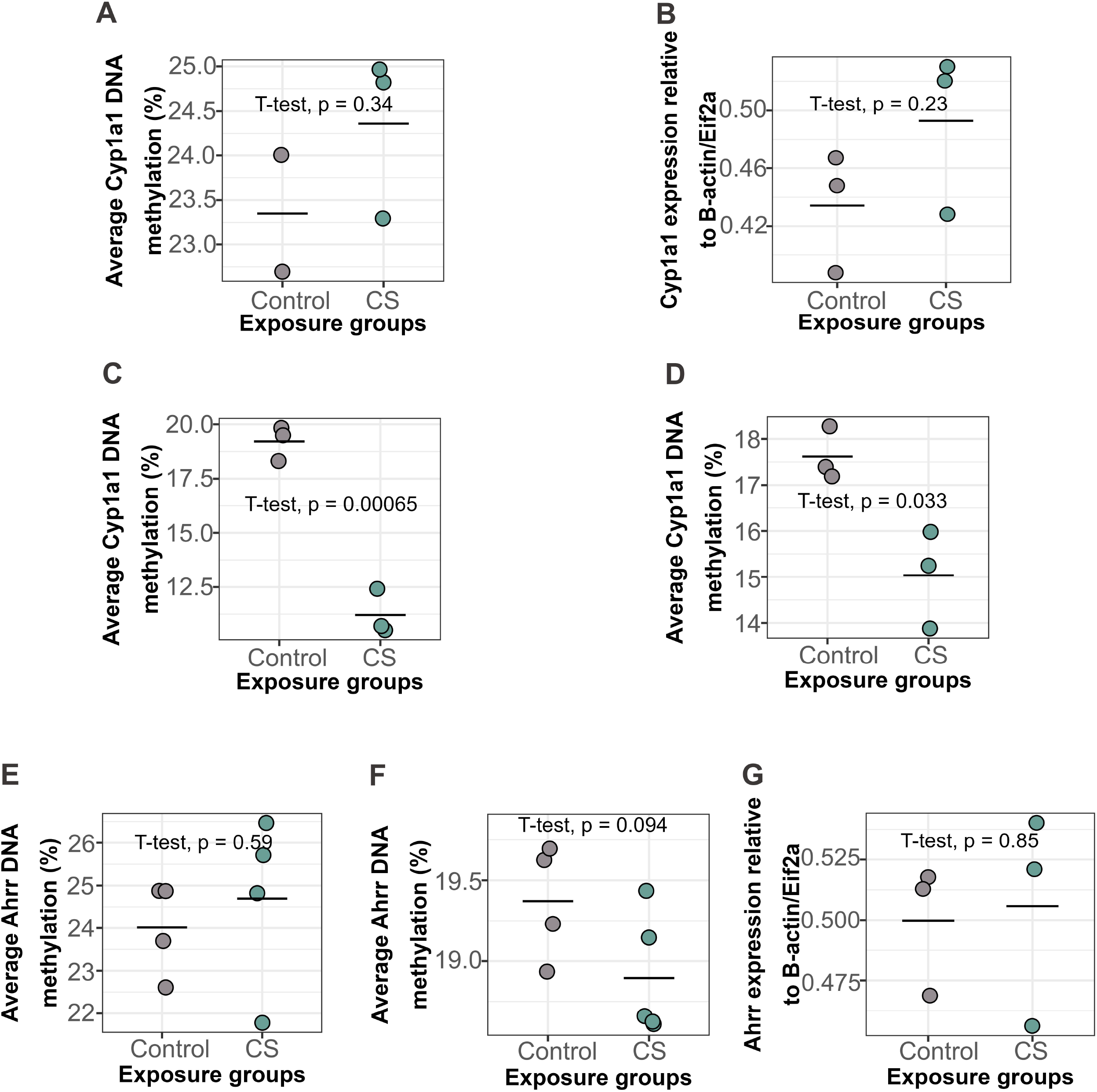
*Cyp1a1* and *Ahrr* DNAm and expression levels in dam blood and lungs immediately after smoking and 15 weeks after smoking cessation. (A) *Cyp1a1* DNAm in dam blood 15 weeks after smoking cessation. (B) *Cyp1a1* expression in dam lungs 15 weeks after smoking cessation. (C) *Cyp1a1* DNAm in dam lungs immediately after 9 weeks of smoking (D) *Cyp1a1* DNAm in dam lungs 15 weeks after smoking cessation. (E) *Ahrr* DNAm in dam blood after 15 weeks of smoking cessation. (F) *Ahrr* DNAm in dam lungs after 15 weeks of smoking cessation. (G) *Ahrr* expression in dam lungs after 15 weeks of smoking cessation. N = 2-5 per group. Differences in DNAm and expression were analyzed using student t-tests.

Conversely, we found no significant differences in *Ahrr* DNAm in dam blood 15 weeks after smoking cessation (Figure 4E). Similarly, *Ahrr* DNAm (Figure 4F) and *Ahrr* expression (Figure 4G) in dam lungs at this same timepoint was also not significantly different between groups.

These results thus show that personal smoking for 9 weeks causes alterations in *Cyp1a1* DNAm in the lungs which persist for 15 weeks after smoking cessation.

### Measurements of effects of smoking on mouse lungs across the whole mouse epigenome using the Illumina mouse methylation microarrays

Following candidate gene DNAm measurements at *Ahrr* and *Cyp1a1*, we evaluated the association between smoking and epigenome-wide DNAm in the lungs immediately after 9 weeks of smoking, using normalized β values from the Illumina mouse microarrays. Due to our small sample size (N = 2-6 per group), we used a combination of *p* value (*p* < 1e-3) and effect size (0.05) cutoffs to determine significance. After adjusting for the 1 surrogate variable recommended by SVA (Supplementary Figure S1A), we performed multivariable regression on beta values from the autosomal probes (Supplementary Figure S1B) and identified 40 significant CpGs differentially methylated in dams exposed to heavy doses of CS for 9 weeks (Table 1). Maternal smoking in pregnancy led to an increase in 37.5% (15 CpGs) and a decrease in 62.5% (25 CpGs) of these significant CpGs (Table 1). While our candidate gene, *Cyp1a1*, was not among our list of significant genes, a closer look at each of the *Cyp1a1* probes on the mouse array reveal large differences in DNAm between controls and CS-exposed dams in 3 of 9 *Cyp1a1* probes that passed preprocessing (Supplementary Figure S2, positions E, F and G). The second candidate gene, *Ahrr,* was also not significant in our epigenome-wide analysis and showed small effect size differences overall (Supplementary Figure S3).

**Table 1:** CpGs significantly altered immediately after exposure to CS for 9 weeks. Linear regression was computed using LIMMA, after adjusting for 1 surrogate variable recommended by SVA (see Figure S1). Results are sorted by CpG name. Direction: direction of change in DNAm compared to controls. Gene: gene names as annotated by UCSC Genome browser. Strand = direction of gene on the chromosomal strand. (“+” = forward, “-” = reverse). Feature = gene region feature as annotated by UCSC Genome browser (tss_body = located within the gene body, tss_200 = located 0-200 bases upstream of transcriptional start site, tss_1500 = located 200-1500 bases upstream of transcriptional start site). N = 3 Control and N = 3 CS. CpG sites were considered significant at *p* < 1e-3 and effect size > 0.05.

We performed a similar analysis on probes from the X chromosome, and after adjusting for the 1 surrogate variable recommended by SVA, found no significantly altered CpGs in dam lungs immediately after smoking (Supplementary Figure S4).

### Gene Ontology terms associated with smoking in the lungs

To gain insights into biological processes potentially affected by smoking in the lungs, we performed gene ontology analyses on genes mapping to the identified 40 significant CpGs. Gene ontology analysis revealed 54 biological processes which passed the FDR cutoff of 0.05 (Table 2). The top pathway findings were mostly driven by *Sox11, Wnt5a, Npnt, Slit3, Magi2* and *Furin*, all being involved in numerous processes. The most significantly implicated pathway with the smallest FDR (FDR = 0.007) was involved in regulation of transmembrane receptor protein serine/threonine kinase signaling pathways.

**Table 2:** List of significantly enriched biological pathways/processes immediately after 9 weeks of smoking. ID: gene ontology identification code. Description: description of the enriched process. Gene ID: genes contributing to enrichment of the biological process. Gene count: number of genes contributing to the biological process. Biological processes were considered significantly enriched at FDR < 0.05.

## DISCUSSION

Cigarette smoke is a complex mixture of over 7000 components^41–44^, which affect virtually all body systems via mechanisms such as inflammation and DNA damage. While the processes linking smoking and adverse health outcomes are poorly understood, DNA methylation alterations have been identified as possible links due to their sensitivity to the environment and relative stability^45^. The association between smoking and changes in DNAm has been well documented, especially in human blood^6–9^. However, while there may be shared DNAm patterns between tissues^14,18^, there is evidence that DNAm can be highly variable across tissues^15–17^, emphasizing the need to study smoke-induced DNAm alterations in more directly affected tissues such as the lungs. The major aim of this study therefore, was to identify DNAm changes in mouse lungs arising due to smoking, and determine how these patterns would change after a period of smoking cessation. To this effect, we adapted an effective smoking mouse model which we have also used to measure the effects of early life smoke exposure on offspring^23^.

In this present study, we observed that the changes in lung function immediately after smoking for 9 weeks were small and not significant. However, 15 weeks after smoking cessation, lung function in CS-exposed dams had significantly declined, as evidenced by increased responsiveness to individual doses of methacholine both in small and large airways. One reason why lung function worsens over time in our smoke-exposed mice may be due to aging. Past research has shown that lung function inevitably declines with age due to structural and immunological factors which impair gas exchange and increase susceptibility to infections^46–48^. However, other studies have reported that smoking exacerbates this decline, even in ex-smokers^49–52^. On the other hand, this trend of worsening/decline in lung function is a marked characteristic of COPD^53,54^. In line with our finding of sustained lung function deficits despite cessation, a large multi-ethnic pooled cohort study^52^, smoking cessation was reported to slightly improve lung function, but these “improvements” resulted in lung function similar to light/low-intensity current smokers. In fact, smoking cessation does not completely restore lung function to the levels of never-smokers^52^. This sustained pattern of lung function decline despite cessation may be due to sustained dysregulation of epigenetic patterns^55–57^, immune responses^58–60^ and airway hyperresponsiveness^61^.

Studies implicating epigenetic alterations as the link between smoking and health outcomes are rapidly growing. Multiple studies have identified changes in DNAm as mediators between smoking and health outcomes both in adult smokers^13,62^ and in offspring exposed to CS *in utero*^23,63–65^. DNAm patterns resulting from smoking may be the key to developing strategies to counteract or prevent its detrimental effects. The fact that our candidate gene analysis did not reveal significant differences in *Ahrr* DNAm or expression in blood or lungs was interesting since the corresponding human locus (cg05575921) is known to be significantly altered in blood of humans exposed to CS^26,66–68^. However, as *Ahrr* is not conserved between mouse and humans, this observation is no completely surprising and further mapping of DNAm in the *Ahrr* gene may reveal deeper insights.

We found that heavy smoking for 9 weeks caused a significant decrease in *Cyp1a1* DNAm immediately after smoking, and that this pattern remained significant till 15 weeks after smoking cessation. However, *Cyp1a1* expression in mouse lungs after smoking cessation, though increased, was not significant, indicating that DNAm changes may last after expression differences have subsided. *Cyp1a1* belongs to the cytochrome p450 enzyme subclass, and is involved in drug/xenobiotic metabolism^69–71^. Therefore, reduced DNAm of *Cyp1a1* in CS-exposed mice as observed in this study could indicate induction of the enzyme to aid detoxification of CS components. These results mimic reduced *Cyp1a1* DNAm and increased expression patterns observed in the lungs^72^, adipose tissue^73^, prostate cancer tissues^74^, buccal cells^19^ and blood^19^ of humans exposed to CS in previous studies.

Prior to this current research, most of the information currently known about smoking and DNAm has been from studies conducted in whole blood in humans. We found three studies to date which directly measured effects of personal smoking on the lungs. The first group measured epigenome-wide DNAm in the blood of current and never-smokers. Next, using the identified significant CpGs from the epigenome-wide analysis, they measured DNAm in the lungs of current, ex- and never-smokers using pyrosequencing^14^. They found a total of 5 of the 15 CpGs in the blood which showed decreased DNAm in the lungs, confirming that both tissues share some similarities but that not all blood-based findings will replicate in the lung^14^. Two of their 5 CpGs mapped to *AHRR*, and one corresponded to the human position we used to identify our mouse candidate *Ahrr* position (cg05575921). In the second study, the effects of a) whole-body CS, b) the tobacco carcinogen 4-(methylnitrosamino)-1-(3-pyridyl)-1-butanone, and c) the inflammatory agent lipopolysaccharide were measured in mouse lungs using RRBS^20^. They found DNAm and hydroxymethylation differences across multiple differentially methylated regions^20^. While there were no overlapping differentially methylated sites between their results and ours, they identified 10 hydroxymethylated sites which exactly corresponded to, or whose isoforms corresponded to 10 of our differentially methylated sites (*Strip1, Slit3, Megf6, Magi2, Atxn2, Sox10/Sox18, Wnt5b, Fkbp3, Asb13* and *Fbxo41/Fbxo45*), effectively corroborating results from our study. However, the disadvantage of RRBS is its poor coverage of intergenic regions and distal regulatory elements ^75,76^. In a design similar to ours, the third group exposed mice to 2 hours of CS twice daily, for 4 weeks consecutively in order to analyze the effect of smoking on lung DNAm. Using liquid hybridization capture-based bisulfite sequencing, they found that smoking altered DNAm patterns at numerous genes, most of which were involved in inflammation and in inflammatory injury in COPD, but it was not possible to determine whether specific findings overlapped with ours^21^. We use the Illumina mouse microarrays to measure DNAm here as it investigates a more diverse range of the genome, including CpG islands, shores, shelves, promoters, enhancers, gene bodies and intergenic regions.

In the epigenome-wide DNAm analysis we conducted, we identified 40 CpGs which were significantly associated with smoking in the lungs. Of these, 8 have previously been linked to smoking: Dopa decarboxylase (*Ddc*) in human blood, saliva or immortalized cell ines^77–79^, Slit guidance ligand 3 (*Slit3*) in zebrafish fin-clips^80^ and human lung adenocarcinoma cell lines and tissues^81^, Wingless-type MMTV integration site family, member 5A (*Wnt5a*) in human bronchial epithelial and lung cancer cells, mouse and human lung tissues^82–84^, Ankyrin repeat and SOCS box-containing 18 (*Asb18*) in human lung tumor samples^85^, Nephronectin (*Npnt*) in human lungs and blood^86^, Membrane associated guanylate kinase, WW and PDZ domain containing 2 (*Magi2*) in human blood and bronchial biopsies^87,88^, Ataxin 2 (*Atxn2*) in human buccal cells and blood^89^ and FK506 binding protein 5 (*Fkbp5*) in human blood plasma^90^, while the other 32 are novel.

Of the 40 significant CpGs identified in this study, two mapped to the promoter of *B3gnt3* (UDP-GlcNAc:betaGal beta-1,3-N-acetylglucosaminyltransferase 3) gene. Both CpGs, which are 32 bp apart, showed significantly increased DNAm in the lungs of dams exposed to CS compared to controls. This gene codes for a type II transmembrane protein which is involved in the biosynthesis of poly-N-acetyllactosamine chains, L-selectin ligand biosynthesis, lymphocyte homing and lymphocyte trafficking and recirculation^91,92^. One study reported increased *B3gnt3* expression levels corresponding to increased immune cell infiltration and correlated with poorer outcomes in patients with lung adenocarcinoma^93^. Other studies have also associated differential *B3gnt3* expression levels with development of many other types of cancers^94–98^. It therefore follows that increased DNAm in our two CpG sites, which are located in a weak promoter, could translate to decreased expression of the gene, consequently resulting in increased tolerance to and reduced sensitivity and response to foreign antigens by T cells.

Another CpG with a large change in DNAm in CS-exposed dams mapped to Ataxin 2 (*Atxn2)* gene. This CpG is located in an intragenic exon and in mouse lungs, is strongly/actively transcribed, showing high levels of H3K36me3. Generally, the function of DNAm at exons varies depending on where the exon is located^99–101^, but most studies have discovered positive correlations between DNAm in intergenic exons and expression levels^99,102^. As our significant CpG is located in an intragenic exon, increased DNAm in smoking dams could signal an increase in *Atxn2* expression which may indicate development of malignancies as *Atxn2* has been shown to modify the risk of pancreatic cancer development among smokers^89^.

We found that heavy smoking resulted in a significant decrease in lung DNAm at Dopa decarboxylase (*Ddc*), a gene encoding a protein which catalyzes the final steps of dopamine and serotonin biosynthesis^103^. Genetic variants in *Ddc,*,especially at the introns^78^, has also been associated with smoking behavior and nicotine dependency^77–79^ in African-American and European-American populations. It therefore follows that alterations at this gene as observed in this research could contribute to nicotine dependency in individuals who smoke. It is however important to note that genetic variants or genes alone do not determine susceptibility to addiction, and that the environment plays a major role^104,105^. In a Finnish twin study, it was discovered that the influence of genetics in adolescent smokers was decreased when the adolescents were highly monitored by their parents^106^. Therefore, in our study for example, mice exposed to smoke which are in stressful environments may, in addition to DNAm alterations and gene expression changes, may display increased susceptibility to nicotine addiction. This emphasizes the need for more research to focus on understanding gene-environment interactions, as it could be crucial in deciphering the mechanism behind addiction.

Gene ontology analysis on our list of significant genes revealed enrichment in multiple biological processes, with the most significant being involved in regulation of transmembrane receptor protein serine/threonine kinase signaling pathways. Components of CS, such as nicotine and the CS-specific carcinogen 4-(methylnitrosamino)-1-(3-pyridyl)-1-butanone, when bound to nicotinic-acetylcholine receptors, activate serine/threonine kinase Akt. Activation of Akt resulted in increased phosphorylation of downstream molecules involved in controlling the cell cycle and protein translation, such as glycogen synthase kinase 3 (GSK-3)^107^. This in turn leads to cancer-like phenotypes such as loss of contact inhibition, desensitization to regulation by growth factors, decreased apoptosis, angiogenesis and increased cellular proliferation^107,108^. Since our current study identified an enrichment for genes involved in regulation of serine/threonine Akt pathways, it could support a predisposition to lung tumorigenesis in smoke-exposed mice.

Even though none of the *Cyp1a1* probes in the mouse methylation microarray was among our 40 significant CpGs, we showed that 3 of the 9 *Cyp1a1* probes have large differences in DNAm. Our inability to detect these positions (and likely some other CpGs) as significant is solely due to our small sample size and hence, low statistical power, which is a limitation in this study. So, while these 3 probes met our strict effect size cutoff of 0.05, they did not meet the *p* value cutoff. Future studies with larger sample sizes would be helpful in generating more precise measurements. Finally, it is worth noting that whole lungs were used in this study, making it impossible to determine whether differences in cell composition are masking some of our findings. However, we took two approaches to mitigate this problem. First, we lavaged the lungs prior to analysis to remove immune cells which we know to be different between CS and control animals, making it unlikely that immune cells are responsible for our observed results. To further support this, we found that *Cyp1a1* DNAm in the blood of adult female control mice (Figure 4A) is higher than *Cyp1a1* DNAm in lungs of control mice (Figure 4D), therefore, infiltration of immune cells from blood to lungs is unlikely to cause the decrease in lung *Cyp1a1* DNAm in the present study. Second, we used surrogate variables to control for variance likely associated with cell type. This approach removes the effects of cell type such that further work will be required to determine which lung cell types exhibit the observed DNAm differences.

In summary, we have shown here that heavy smoking alters DNAm patterns in mouse lungs at multiple CpGs implicated in tumorigenesis and metastasis, neurodegenerative disorders, nicotine dependency, immune cell response and lung function defects, at least some of which last up to 15 weeks after smoking cessation. The fact that 75% of the significant CpGs reported here have never been linked to CS in the past, even in studies which investigated effects of CS in the blood, emphasizes the need for more studies to be done on the lungs, as it clearly has different response than other tissues. This research is one part of a much larger study to investigate the effects of early life CS exposure on offspring DNAm patterns. Therefore, the results presented here will hopefully also shed more light on some of the patterns we observe in offspring exposed to CS *in utero*.

**Supplementary Figure S1:**
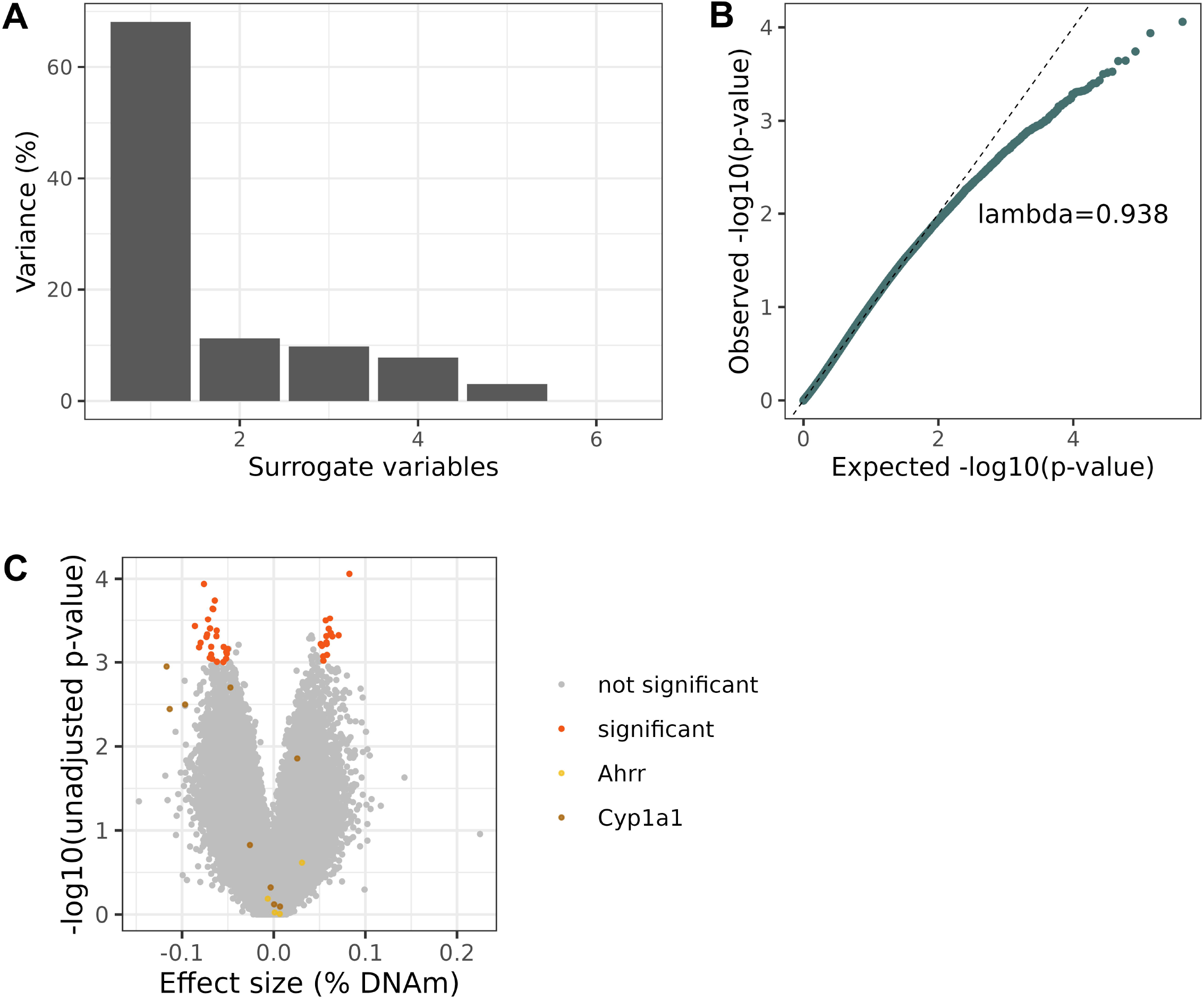
Linear regression models were adequately fit to measure effects of heavy smoking on dam lung DNAm in autosomal probes. N = 3 per group. (A) Surrogate Variable Analysis (SVA) identified 5 technical variables in dam lungs immediately after 9 weeks of heavy smoking. We included the first variable only to our regression model based on SVA recommendations. (B) QQ-plot showed a slightly deflated but adequate fit in our limma model, upon inclusion of the surrogate variable. (C) Volcano plot showing the 40 differentially methylated CpGs (highlighted in orange) and non-significant CpGs (grey). DNAm differences were measured using multivariable linear regression, from the limma package in R. Significance was determined using value cutoff of *p* < 1e-3 and effect size > 5%.

**Supplementary Figure S2:**
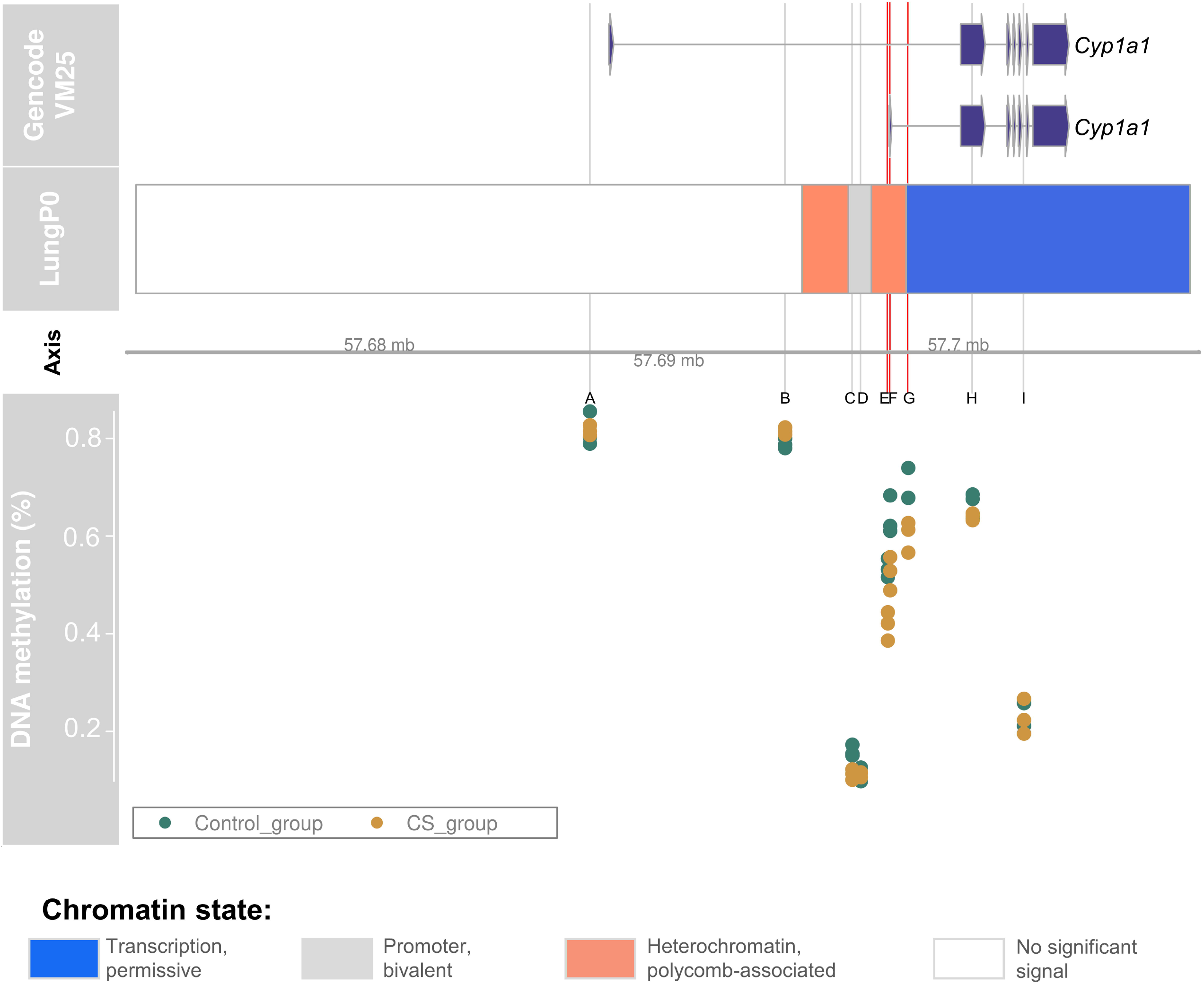
Chromatin state, genomic locations and DNAm levels at 9 *Cyp1a1* positions on the Illumina mouse methylation microarray. *Cyp1a1* sites from the Illumina mouse array are labelled A-I, the 3 sites with the greatest effect sizes are highlighted in red (sites E, F and G), while the other 6 are highlighted in grey. The genomic locations of the 9 *Cyp1a1* array sites are as follows: A: 57687270, B: 57694012, C: 57696335, D: 57696629, E: 57697560, F: 57697641, G: 57698266, H: 57700489 and I: 57702263. Our candidate CpG is at position 57696231. Information on chromatin state was obtained from UCSC genome browser using mouse genome build GRCm38/mm10. Chromatin state displayed here is from mouse lungs at postnatal day 0, as it is the latest timepoint currently available.

**Supplementary Figure S3:**
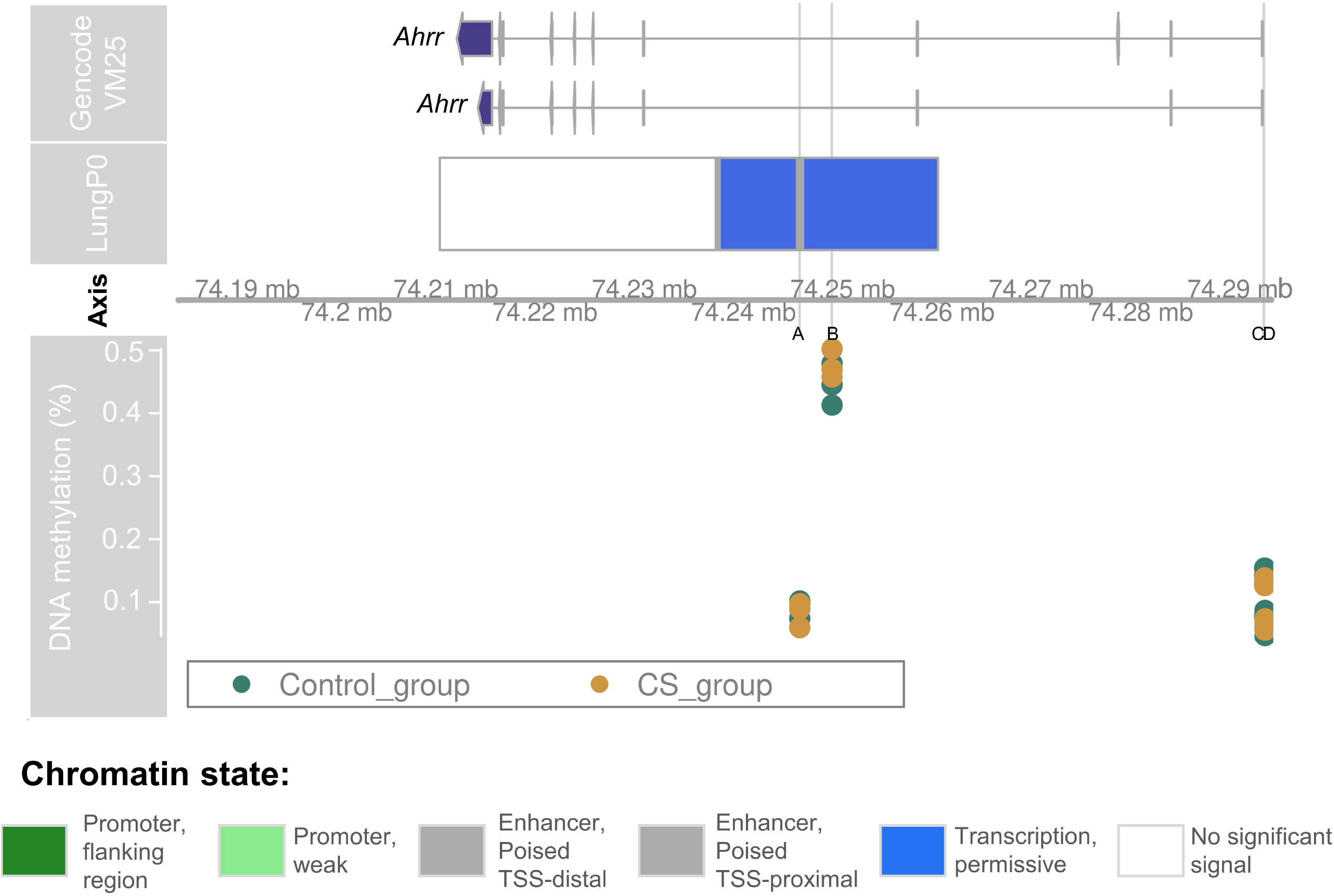
Chromatin state, genomic locations and DNAm levels at 4 *Ahrr* positions on the Illumina mouse methylation microarray. *Ahrr* sites from the Illumina mouse array are labelled A-D and highlighted in grey. The genomic locations of the 4 *Ahrr* array sites are as follows: A: 74245665, B: 74248917, C: 74292502, D: 74292541. Our candidate CpG is at position 74260517. Information on chromatin state was obtained from UCSC genome browser using mouse genome build GRCm38/mm10. Chromatin state displayed here is from mouse lungs at postnatal day 0, as it is the latest timepoint currently available.

**Supplementary Figure S4:**
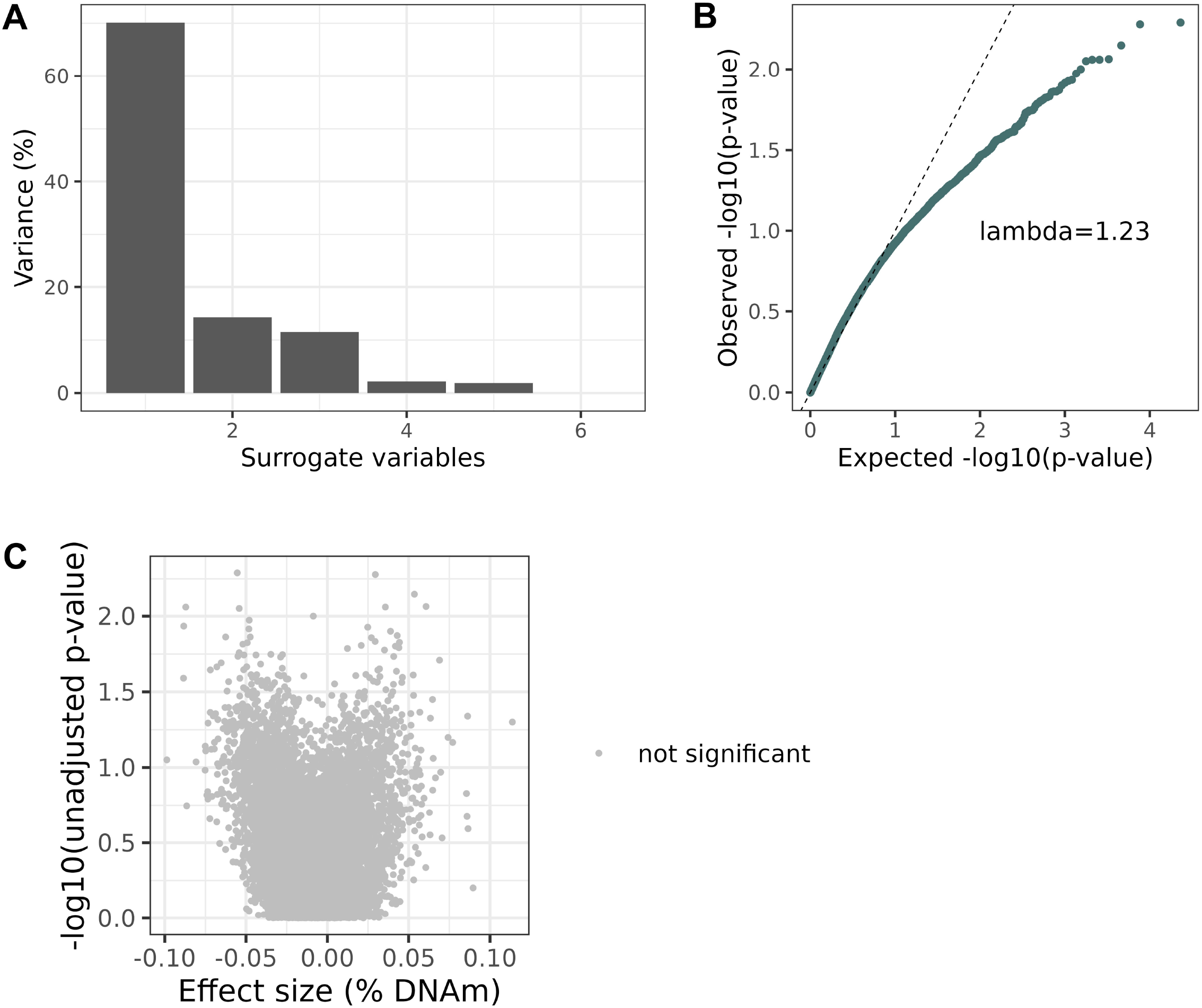
Linear regression models to measure effects of heavy smoking on dam lung DNAm in female X chromosome probes. N = 3 per group. (A) Surrogate Variable Analysis (SVA) identified 5 technical variables in dam lungs immediately after 9 weeks of heavy smoking. We included the first variable only to our regression model based on SVA recommendations. (B) QQ-plot showed a slightly deflated fit in our limma model, upon inclusion of the surrogate variable. (C) Volcano plot showing no significantly differentially methylated CpGs. DNAm differences were measured using multivariable linear regression, from the limma package in R. Significance was determined using value cutoff of *p* < 1e-3 and effect size > 5%.

## REFERENCES

1. Tamimi, A., Serdarevic, D. & Hanania, N. A. The effects of cigarette smoke on airway inflammation in asthma and COPD: Therapeutic implications. Respiratory Medicine 106, 319–328 (2012).

2. Boulet, L.-P. et al. Smoking and Asthma: Clinical and Radiologic Features, Lung Function, and Airway Inflammation. Chest 129, 661–668 (2006).

3. Nakano, Y. et al. Computed Tomographic Measurements of Airway Dimensions and Emphysema in Smokers. Am J Respir Crit Care Med 162, 1102–1108 (2000).

4. Pietinalho, A., Pelkonen, A. & Rytilä, P. Linkage between smoking and asthma. Allergy 64, 1722–1727 (2009).

5. Piipari, R., Jaakkola, J. J. K., Jaakkola, N. & Jaakkola, M. S. Smoking and asthma in adults. European Respiratory Journal 24, 734–739 (2004).

6. Elliott, H. R. et al. Differences in smoking associated DNA methylation patterns in South Asians and Europeans. Clinical epigenetics 6, 4 (2014).

7. Su, D. et al. Distinct Epigenetic Effects of Tobacco Smoking in Whole Blood and among Leukocyte Subtypes. PLOS ONE 11, e0166486 (2016).

8. Zhang, Y., Florath, I., Saum, K.-U. & Brenner, H. Self-reported smoking, serum cotinine, and blood DNA methylation. Environ Res 146, 395–403 (2016).

9. Zeilinger, S. et al. Tobacco smoking leads to extensive genome-wide changes in DNA methylation. PLoS One 8, e63812 (2013).

10. Monick, M. M. et al. Coordinated changes in AHRR methylation in lymphoblasts and pulmonary macrophages from smokers. American journal of medical genetics. Part B, Neuropsychiatric geneticsLJ: the official publication of the International Society of Psychiatric Genetics 159B, 141–151 (2012).

11. Oliveira, N. F. P. et al. DNA methylation status of the IL8 gene promoter in oral cells of smokers and non-smokers with chronic periodontitis. Journal of Clinical Periodontology 36, 719–725 (2009).

12. Bollepalli, S., Korhonen, T., Kaprio, J., Anders, S. & Ollikainen, M. EpiSmokEr: a robust classifier to determine smoking status from DNA methylation data. Epigenomics 11, 1469–1486 (2019).

13. Zhang, Y. et al. Smoking-associated DNA methylation markers predict lung cancer incidence. Clinical Epigenetics 8, 127 (2016).

14. de Vries, M. et al. From blood to lung tissue: effect of cigarette smoke on DNA methylation and lung function. Respir. Res. 19, 212 (2018).

15. Bloushtain-Qimron, N. et al. Cell type-specific DNA methylation patterns in the human breast. Proceedings of the National Academy of Sciences 105, 14076–14081 (2008).

16. Zhou, J. et al. Tissue-specific DNA methylation is conserved across human, mouse, and rat, and driven by primary sequence conservation. BMC Genomics 18, 724 (2017).

17. Scherer, M. et al. Identification of tissue-specific and common methylation quantitative trait loci in healthy individuals using MAGAR. Epigenetics & Chromatin 14, 44 (2021).

18. Stueve, T. R. et al. Epigenome-wide analysis of DNA methylation in lung tissue shows concordance with blood studies and identifies tobacco smoke-inducible enhancers. Human Molecular Genetics 26, 3014–3027 (2017).

19. Teschendorff, A. E. et al. Correlation of Smoking-Associated DNA Methylation Changes in Buccal Cells With DNA Methylation Changes in Epithelial Cancer. JAMA Oncology 1, 476–485 (2015).

20. Seiler, C. L. et al. Inhalation exposure to cigarette smoke and inflammatory agents induces epigenetic changes in the lung. Sci Rep 10, 11290 (2020).

21. Li, P. et al. DNA Methylation Profiling in a Cigarette Smoke-Exposed Mouse Model of Airway Inflammation. International Journal of Chronic Obstructive Pulmonary Disease 17, 2443–2450 (2022).

22. Zhou, W. et al. DNA methylation dynamics and dysregulation delineated by high-throughput profiling in the mouse. Cell Genomics 2, 100144 (2022).

23. Onuzulu, C. D. et al. Early life exposure to cigarette smoke primes lung function and DNA methylation changes at Cyp1a1 upon exposure later in life. American Journal of Physiology-Lung Cellular and Molecular Physiology (2023) doi:10.1152/ajplung.00192.2023.

24. Ma, Y. et al. Sustained suppression of IL-13 by a vaccine attenuates airway inflammation and remodeling in mice. Am J Respir Cell Mol Biol 48, 540–549 (2013).

25. Ryu, M. H. et al. Chronic exposure to perfluorinated compounds: Impact on airway hyperresponsiveness and inflammation. Am J Physiol Lung Cell Mol Physiol 307, L765–774 (2014).

26. Joubert, B. R. et al. DNA Methylation in Newborns and Maternal Smoking in Pregnancy: Genome-wide Consortium Meta-analysis. American journal of human genetics 98, 680– 696 (2016).

27. MUSCLE: multiple sequence alignment with improved accuracy and speed | IEEE Conference Publication | IEEE Xplore. https://ieeexplore.ieee.org/abstract/document/1332560.

28. Livak, K. J. & Schmittgen, T. D. Analysis of relative gene expression data using real-time quantitative PCR and the 2(-Delta Delta C(T)) Method. Methods 25, 402–408 (2001).

29. Wang, R. et al. Long non11coding RNA HOX transcript antisense RNA promotes expression of 14113113σ in non11small cell lung cancer. Experimental and Therapeutic Medicine 14, 4503–4508 (2017).

30. Zhou, W., Triche, T. J., Laird, P. W. & Shen, H. SeSAMe: reducing artifactual detection of DNA methylation by Infinium BeadChips in genomic deletions. Nucleic Acids Research 19, 129 (2018).

31. Triche, T. J., Weisenberger, D. J., Van Den Berg, D., Laird, P. W. & Siegmund, K. D. Low-level processing of Illumina Infinium DNA Methylation BeadArrays. Nucleic Acids Research 41, e90–e90 (2013).

32. Johnson, W. E., Li, C. & Rabinovic, A. Adjusting batch effects in microarray expression data using empirical Bayes methods. Biostatistics 8, 118–127 (2007).

33. Leek, J. T. & Storey, J. D. Capturing Heterogeneity in Gene Expression Studies by Surrogate Variable Analysis. PLOS Genetics 3, e161 (2007).

34. Gómez-de-Mariscal, E. et al. Use of the p-values as a size-dependent function to address practical differences when analyzing large datasets. Sci Rep 11, 20942 (2021).

35. Wu, T. et al. clusterProfiler 4.0: A universal enrichment tool for interpreting omics data. The Innovation 2, 100141 (2021).

36. Leong, J. W. et al. The Elimination Half-Life of Urinary Cotinine in Children of Tobacco-Smoking Mothers. Pulmonary Pharmacology & Therapeutics 11, 287–290 (1998).

37. Benowitz, N. L. Cotinine as a Biomarker of Environmental Tobacco Smoke Exposure. Epidemiologic Reviews 18, 188–204 (1996).

38. Melgert, B. N. et al. Short-Term Smoke Exposure Attenuates Ovalbumin-Induced Airway Inflammation in Allergic Mice. Am J Respir Cell Mol Biol 30, 880–885 (2004).

39. Blacquière, M. J. et al. Maternal smoking during pregnancy induces airway remodelling in mice offspring. European Respiratory Journal 33, 1133–1140 (2009).

40. Lei, Y., Cao, Y.-X., Xu, C.-B. & Zhang, Y. The Raf-1 inhibitor GW5074 and dexamethasone suppress sidestream smoke-induced airway hyperresponsiveness in mice. Respir Res 9, 71 (2008).

41. Colombo, G. et al. Oxidative damage in human gingival fibroblasts exposed to cigarette smoke. Free Radical Biology and Medicine 52, 1584–1596 (2012).

42. Zeng, T. et al. Advanced Materials Design for Adsorption of Toxic Substances in Cigarette Smoke. Advanced Science 10, 2301834 (2023).

43. Soleimani, F., Dobaradaran, S., De-la-Torre, G. E., Schmidt, T. C. & Saeedi, R. Content of toxic components of cigarette, cigarette smoke vs cigarette butts: A comprehensive systematic review. Science of The Total Environment 813, 152667 (2022).

44. Staal, Y. C. M., Bos, P. M. J. & Talhout, R. Methodological Approaches for Risk Assessment of Tobacco and Related Products. Toxics 10, 491 (2022).

45. Nwanaji-Enwerem, J. C. & Colicino, E. DNA Methylation–Based Biomarkers of Environmental Exposures for Human Population Studies. Curr Envir Health Rpt 7, 121– 128 (2020).

46. Thomas, E. T., Guppy, M., Straus, S. E., Bell, K. J. L. & Glasziou, P. Rate of normal lung function decline in ageing adults: a systematic review of prospective cohort studies. BMJ Open 9, e028150 (2019).

47. Schneider, J. L. et al. The aging lung: Physiology, disease, and immunity. Cell 184, 1990– 2019 (2021).

48. Sharma, G. & Goodwin, J. Effect of aging on respiratory system physiology and immunology. Clin Interv Aging 1, 253–260 (2006).

49. Scanlon, P. D. et al. Smoking Cessation and Lung Function in Mild-to-Moderate Chronic Obstructive Pulmonary Disease. Am J Respir Crit Care Med 161, 381–390 (2000).

50. Burchfiel, C. M. et al. Effects of smoking and smoking cessation on longitudinal decline in pulmonary function. Am J Respir Crit Care Med 151, 1778–1785 (1995).

51. Bossé, R., Sparrow, D., Rose, C. L. & Weiss, S. T. Longitudinal Effect of Age and Smoking Cessation on Pulmonary Function. Am Rev Respir Dis 123, 378–381 (1981).

52. Oelsner, E. C. et al. Lung function decline in former smokers and low-intensity current smokers: the NHLBI Pooled Cohorts Study. Lancet Respir Med 8, 34–44 (2020).

53. Donaldson, G. C. et al. Airway and Systemic Inflammation and Decline in Lung Function in Patients With COPD. Chest 128, 1995–2004 (2005).

54. Donaldson, G. C., Seemungal, T. a. R., Bhowmik, A. & Wedzicha, J. A. Relationship between exacerbation frequency and lung function decline in chronic obstructive pulmonary disease. Thorax 57, 847–852 (2002).

55. Soria, J.-C. et al. Aberrant promoter methylation of multiple genes in bronchial brush samples from former cigarette smokers. Cancer Res 62, 351–355 (2002).

56. Belinsky, S. A. et al. Aberrant promoter methylation in bronchial epithelium and sputum from current and former smokers. Cancer Res 62, 2370–2377 (2002).

57. Wang, G. et al. Persistence of Smoking-Induced Dysregulation of MiRNA Expression in the Small Airway Epithelium Despite Smoking Cessation. PLoS One 10, e0120824 (2015).

58. Hogg, J. C. Why does airway inflammation persist after the smoking stops? Thorax 61, 96 (2006).

59. Rutgers, S. et al. Ongoing airway inflammation in patients with COPD who do not currently smoke. Thorax 55, 12 (2000).

60. Shiels, M. S. et al. Cigarette Smoking and Variations in Systemic Immune and Inflammation Markers. JNCI Journal of the National Cancer Institute 106, (2014).

61. Lapperre, T. S. et al. Relation between duration of smoking cessation and bronchial inflammation in COPD. Thorax 61, 115 (2006).

62. Wang, T., Xia, P. & Su, P. High-Dimensional DNA Methylation Mediates the Effect of Smoking on Crohn’s Disease. Frontiers in Genetics 13, (2022).

63. Xu, R. et al. DNA methylation mediates the effect of maternal smoking on offspring birthweight: a birth cohort study of multi-ethnic US mother–newborn pairs. Clin Epigenet 13, 47 (2021).

64. Wiklund, P. et al. DNA methylation links prenatal smoking exposure to later life health outcomes in offspring. Clinical Epigenetics 11, 97 (2019).

65. Hannon, E. et al. Variable DNA methylation in neonates mediates the association between prenatal smoking and birth weight. Philosophical Transactions of the Royal Society B: Biological Sciences 374, 20180120 (2019).

66. Joubert, B. R. et al. 450K epigenome-wide scan identifies differential DNA methylation in newborns related to maternal smoking during pregnancy. Environmental health perspectives 120, 1425–1431 (2012).

67. Lee, K. W. K. et al. Prenatal Exposure to Maternal Cigarette Smoking and DNA Methylation: Epigenome-Wide Association in a Discovery Sample of Adolescents and Replication in an Independent Cohort at Birth through 17 Years of Age. Environ Health Perspect 123, 193–199 (2015).

68. Richmond, R. C. et al. Prenatal exposure to maternal smoking and offspring DNA methylation across the lifecourse: findings from the Avon Longitudinal Study of Parents and Children (ALSPAC). Hum Mol Genet 24, 2201–2217 (2015).

69. Ma, Q. Induction of CYP1A1. The AhR / DRE Paradigm Transcription, Receptor Regulation, and Expanding Biological Roles. Current Drug Metabolism 2, 149–164 (2001).

70. Nebert, D. W. et al. Role of the aromatic hydrocarbon receptor and [Ah] gene battery in the oxidative stress response, cell cycle control, and apoptosis. Biochem Pharmacol 59, 65–85 (2000).

71. Ohashi, H. et al. The aryl hydrocarbon receptor–cytochrome P450 1A1 pathway controls lipid accumulation and enhances the permissiveness for hepatitis C virus assembly. J Biol Chem 293, 19559–19571 (2018).

72. Tekpli, X. et al. DNA methylation of the CYP1A1 enhancer is associated with smoking-induced genetic alterations in human lung. International Journal of Cancer 131, 1509– 1516 (2012).

73. Tsai, P.-C. et al. Smoking induces coordinated DNA methylation and gene expression changes in adipose tissue with consequences for metabolic health. Clin Epigenet 10, 126 (2018).

74. Mitsui, Y. et al. Functional role and tobacco smoking effects on methylation of CYP1A1 gene in prostate cancer. Oncotarget 7, 49107 (2016).

75. Yong, W.-S., Hsu, F.-M. & Chen, P.-Y. Profiling genome-wide DNA methylation. Epigenetics & Chromatin 9, 26 (2016).

76. Rauluseviciute, I., Drabløs, F. & Rye, M. B. DNA methylation data by sequencing: experimental approaches and recommendations for tools and pipelines for data analysis. Clinical Epigenetics 11, 193 (2019).

77. Ma, J. Z. et al. Haplotype analysis indicates an association between the DOPA decarboxylase (DDC) gene and nicotine dependence. Hum Mol Genet 14, 1691–1698 (2005).

78. Yu, Y. et al. Intronic variants in the dopa decarboxylase (DDC) gene are associated with smoking behavior in European-Americans and African-Americans. Hum Mol Genet 15, 2192–2199 (2006).

79. Zhang, H. et al. DOPA decarboxylase gene is associated with nicotine dependence. Pharmacogenomics 7, 1159–1166 (2006).

80. García-González, J. et al. Identification of slit3 as a locus affecting nicotine preference in zebrafish and human smoking behaviour. eLife 9, e51295.

81. Wang, J. et al. Expression and prognosis effect of methylation-regulated SLIT3 and SPARCL1 genes in smoking-related lung adenocarcinoma. Zhonghua yi xue za zhi 99, 1553–1557 (2019).

82. Whang, Y. M. et al. Wnt5a is associated with cigarette smoke-related lung carcinogenesis via protein kinase C. PLoS One 8, e53012 (2013).

83. Hussain, M. et al. Tobacco smoke induces polycomb-mediated repression of Dickkopf-1 in lung cancer cells. Cancer Res 69, 3570–3578 (2009).

84. Feller, D. et al. Cigarette Smoke-Induced Pulmonary Inflammation Becomes Systemic by Circulating Extracellular Vesicles Containing Wnt5a and Inflammatory Cytokines. Front Immunol 9, 1724 (2018).

85. Shen, Z. et al. Potential Genes Associated with the Survival of Lung Adenocarcinoma Were Identified by Methylation. Computational and Mathematical Methods in Medicine 2020, 1–13 (2020).

86. Wain, L. V. et al. Novel insights into the genetics of smoking behaviour, lung function, and chronic obstructive pulmonary disease (UK BiLEVE): a genetic association study in UK Biobank. Lancet Respir Med 3, 769–781 (2015).

87. Quach, B. C. et al. Expanding the genetic architecture of nicotine dependence and its shared genetics with multiple traits. Nat Commun 11, 5562 (2020).

88. Dijkstra, A. E. et al. Novel Genes for Airway Wall Thickness Identified with Combined Genome-Wide Association and Expression Analyses. Am J Respir Crit Care Med 191, 547–556 (2015).

89. Tang, H. et al. Axonal guidance signaling pathway interacting with smoking in modifying the risk of pancreatic cancer: a gene- and pathway-based interaction analysis of GWAS data. Carcinogenesis 35, 1039–1045 (2014).

90. Koopmann, A. et al. The Effect of Nicotine on HPA Axis Activity in Females is Modulated by the FKBP5 Genotype. Annals of Human Genetics 80, 154–161 (2016).

91. Yeh, J. C. et al. Novel sulfated lymphocyte homing receptors and their control by a Core1 extension beta 1,3-N-acetylglucosaminyltransferase. Cell 105, 957–969 (2001).

92. Hennet, T. et al. Genomic Cloning and Expression of Three Murine UDP-galactose: β-N-Acetylglucosamine β1,3-Galactosyltransferase Genes *. Journal of Biological Chemistry 273, 58–65 (1998).

93. Leng, X. et al. Identifying the prognostic significance of B3GNT3 with PD-L1 expression in lung adenocarcinoma. Transl Lung Cancer Res 10, 965–980 (2021).

94. Gupta, R. et al. Global analysis of human glycosyltransferases reveals novel targets for pancreatic cancer pathogenesis. Br J Cancer 122, 1661–1672 (2020).

95. Zhuang, H. et al. B3GNT3 overexpression promotes tumor progression and inhibits infiltration of CD8+ T cells in pancreatic cancer. Aging (Albany NY) 13, 2310–2329 (2020).

96. Xu, J., Guo, Z., Yuan, S., Li, H. & Luo, S. Upregulation of B3GNT3 is associated with immune infiltration and activation of NF-κB pathway in gynecologic cancers. J Reprod Immunol 152, 103658 (2022).

97. Lu, J. et al. The role of B3GNT3 as an oncogene in the growth, invasion and migration of esophageal cancer cells. Acta Cir Bras 38, e380923 (2023).

98. Zhou, H., Zhao, J., Yang, X., Liu, J. & Huang, W. Study on the Expression of β-1,3-N-acetylglucosaminyltransferase 3 in Gastric Cancer and the Mechanism Promoting Gastric Cancer Progression Based on the Extraction Method of Nanomagnetic Beads. J Biomed Nanotechnol 18, 677–692 (2022).

99. Li, S., Zhang, J., Huang, S. & He, X. Genome-wide analysis reveals that exon methylation facilitates its selective usage in the human transcriptome. Brief Bioinform 19, 754–764 (2018).

100. Song, K., Li, L. & Zhang, G. The association between DNA methylation and exon expression in the Pacific oyster Crassostrea gigas. PLoS One 12, e0185224 (2017).

101. Brenet, F. et al. DNA Methylation of the First Exon Is Tightly Linked to Transcriptional Silencing. PLOS ONE 6, e14524 (2011).

102. Chuang, T.-J., Chen, F.-C. & Chen, Y.-Z. Position-dependent correlations between DNA methylation and the evolutionary rates of mammalian coding exons. Proc Natl Acad Sci U S A 109, 15841–15846 (2012).

103. Ng, J., Papandreou, A., Heales, S. J. & Kurian, M. A. Monoamine neurotransmitter disorders—clinical advances and future perspectives. Nat Rev Neurol 11, 567–584 (2015).

104. Kirst, M., Mecredy, G., Borland, T. & Chaiton, M. Predictors of Substance Use Among Young Adults Transitioning Away from High School: A Narrative Review. Substance Use & Misuse 49, 1795–1807 (2014).

105. Vink, J. M. Genetics of Addiction: Future Focus on Gene × Environment Interaction? J. Stud. Alcohol Drugs 77, 684–687 (2016).

106. Dick, D. M. et al. Parental monitoring moderates the importance of genetic and environmental influences on adolescent smoking. Journal of Abnormal Psychology 116, 213–218 (2007).

107. West, K. A. et al. Rapid Akt activation by nicotine and a tobacco carcinogen modulates the phenotype of normal human airway epithelial cells. J Clin Invest 111, 81–90 (2003).

108. Schuller, H. M. Mechanisms of smoking-related lung and pancreatic adenocarcinoma development. Nat Rev Cancer 2, 455–463 (2002).

